# A MYC Family Switch: L-MYC Drives and Maintains Neuroendocrine Lineage Programs in Prostate Cancer

**DOI:** 10.64898/2025.12.25.696507

**Authors:** Jeyaluxmy Sivalingam, Kyung Hyun Cho, Yingli Shi, Preston Barron, Michael Lan, Omar Franco, Xiuping Yu

**Affiliations:** Department of Biochemistry & Molecular Biology, LSU Health-Shreveport, LA; Department of Microbiology & Immunology, LSU Health-Shreveport, LA; School of Medicine, LSU Health-Shreveport, LA; Department of Genetics, LSU Health-New Orleans, LA; Department of Urology, LSU Health-Shreveport, LA

## Abstract

Neuroendocrine prostate cancer (NEPC) is an aggressive, therapy-resistant subtype that emerges through lineage plasticity following androgen receptor (AR) pathway inhibition. Although MYC family oncogenes are central to prostate cancer progression, the role of MYCL (L-MYC) in NEPC has remained unclear. Here, we show that MYCL is selectively and robustly upregulated in NEPC patient samples and cell line models, whereas MYC is downregulated and MYCN remains low, revealing a lineage-associated MYC family switch. MYCL expression strongly correlates with the neuroendocrine lineage regulators ASCL1 and INSM1 and inversely with adenocarcinoma-associated genes. Mechanistically, MYCL activation is not driven by genomic amplification but reflects a permissive epigenetic landscape. Functionally, MYCL overexpression suppresses AR signaling and induces neuroendocrine-like transcriptional reprogramming, whereas MYCL knockdown disrupts neuroendocrine lineage identity and restores adenocarcinoma-associated gene expression, including MYC. We further identify ASCL1 and INSM1 as upstream regulators of MYCL, establishing a conserved neuroendocrine transcriptional axis. Together, these findings define MYCL as a lineage-specific regulator that drives neuroendocrine identity and plasticity in advanced prostate cancer.

## INTRODUCTION

Prostate cancer (PCa) is the second most diagnosed nonskin cancer and the fifth leading cause of cancer-related deaths in men worldwide. In the United States, it ranks first in incidence and second in cancer related death (Rawla, 2019)(Rawla, 2019). Early stage PCa are typically androgen-dependent and respond to androgen deprivation therapy. However, over time, most tumors eventually progress to a castration-resistant state (CRPC), driven by genetic and epigenetic alterations that reactivate androgen receptor (AR) signaling even under low-androgen conditions (Kobayashi et al., 2013)(Kobayashi et al., 2013).

While many CRPC cases continue to respond to AR pathway inhibitors (ARPIs), a subset of tumors becomes resistant through lineage plasticity, leading to the emergence of neuroendocrine prostate cancer (NEPC). The incidence of NEPC has increased since the widespread clinical use of next generation of ARPIs around 2012. Notably, NEPC and adenocarcinoma prostate cancer (AdPC) share a largely conserved genomic landscape, but exhibit distinct epigenomic signatures (Beltran et al., 2016)(Beltran et al., 2016). During this phenotypic shift, AR expression is lost and neuroendocrine (NE) lineage markers such as ASCL1, INSM1 and CHGA become upregulated (Beltran et al., 2016)(Beltran et al., 2016). Currently, there are no effective treatments for NEPC, and the average survival time remains less than seven months (Zhu et al., 2021)(Zhu et al., 2021). Therefore, a deeper understanding of the molecular mechanisms driving this transdifferentiation continues to be a priority for advancing PCa research.

The Myelocytomatosis (MYC) family of oncogenes, comprising MYC(c-MYC), MYCN(N-MYC), and MYCL(L-MYC), encodes transcription factors that regulate approximately 15% of all genes, including those involved in stress responses, proliferation, differentiation, cell cycle progression, apoptosis, and immune regulation (Ahmadi et al., 2021)(Ahmadi et al., 2021)(Ahmadi et al., 2021)(Ahmadi et al., 2021). The dysregulation of MYC family members has been implicated in tumorigenesis across multiple cancer types, including PCa. In AdPC, MYC is frequently amplified and overexpressed, where it promotes tumor progression(Qiu et al., 2022a)(Qiu et al., 2022a)(Qiu et al., 2022a)(Qiu et al., 2022a), (Taylor et al., 2010)(Taylor et al., 2010)(Taylor et al., 2010)(Taylor et al., 2010), In contrast, MYCN amplification is observed in approximately 40% of NEPC cases (Beltran et al., 2011)(Beltran et al., 2011)(Beltran et al., 2011)(Beltran et al., 2011) Notably, MYCN, in cooperation with activated AKT1, can transform human prostate epithelial cells and drive NEPC progression (Lee et al., 2016)(Lee et al., 2016)(Lee et al., 2016)(Lee et al., 2016). Additionally, MYCN can recruit EZH2 to suppress AR signaling and promote NE features (Dardenne et al., 2016)(Dardenne et al., 2016)(Dardenne et al., 2016)(Dardenne et al., 2016).

While MYC and MYCN have been extensively studied in PCa, MYCL has primarily been described only at the level of focal copy-number amplification in clinically localized AdPC(Boutros et al., 2015)(Boutros et al., 2015)(Boutros et al., 2015)(Boutros et al., 2015). Its regulation and functional role remain poorly understood, especially in NEPC, where MYCL expression is highest (Sivalingam & Yu, 2025)(Sivalingam & Yu, 2025)(Sivalingam & Yu, 2025)(Sivalingam & Yu, 2025).

Our analysis of publicly available patient and cell-line RNA-seq datasets revealed that MYCL expression is selectively upregulated in NEPC, whereas MYC is preferentially enriched in AdPC, and MYCN expression remains comparatively low in NEPC(Sivalingam & Yu, 2025) (Sivalingam & Yu, 2025)(Sivalingam & Yu, 2025) (Sivalingam & Yu, 2025). This reciprocal expression pattern between MYCL and MYC suggests a potential MYC family switch associated with lineage plasticity and neuroendocrine transdifferentiation. Here, we provide the first mechanistic insights into MYCL function in PCa, uncovering its epigenetic regulation and a previously unrecognized role in neuroendocrine lineage maintenance. Together, our findings support MYCL as a key transcriptional regulator required for establishing and maintaining neuroendocrine identity in NEPC.

## RESULTS

### MYCL is upregulated and MYC is downregulated in NEPC across patient samples and cell line models

Transcriptomic analysis of patient cohorts (Beltran, ProAtlas) and prostate cancer cell-line models (CTPC) representing adenocarcinoma and neuroendocrine states revealed distinct MYC family expression patterns associated with disease progression. MYC was highly expressed in adenocarcinoma but markedly reduced in NEPC, with MYC protein undetectable in the NCI-H660 NEPC cell line (Fig. 1A-H). In contrast, MYCL showed robust upregulation in NEPC patient samples and was strongly enriched at both the mRNA and protein levels in NCI-H660 cells. MYCN exhibited minimal mRNA expression across patient subtypes and remained undetectable at the protein level in NCI-H660. Consistent with these findings, single-cell RNA-seq analysis of human prostate tumors revealed MYCL enrichment in NEPC as well as in the AR-high adenocarcinoma (AdPC_ARhi) population, a castration-resistant adenocarcinoma state, suggesting that MYCL activation may occur early during neuroendocrine lineage transition (Supplementary Fig. 1A).

**Fig. 1:**
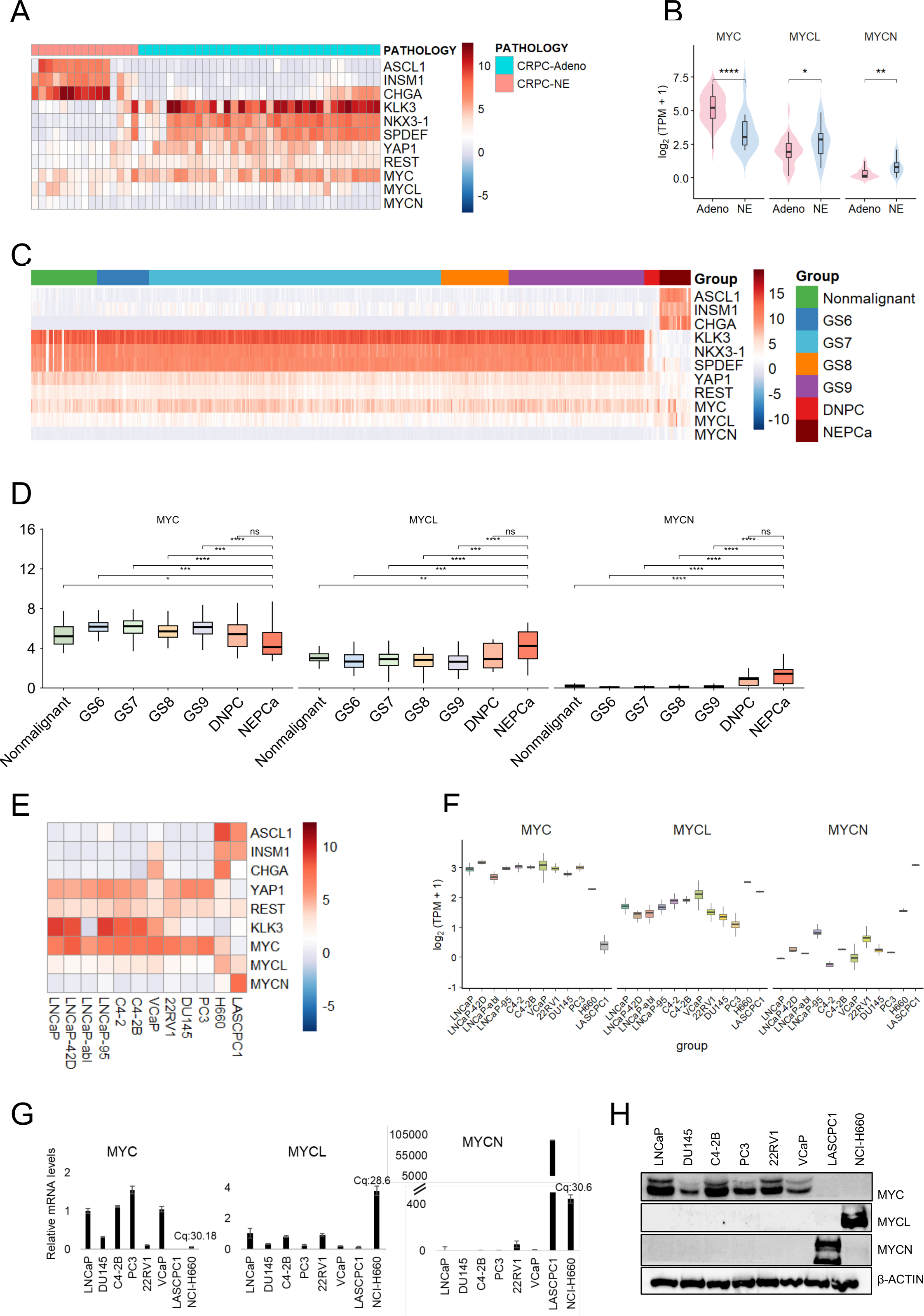
MYCL is upregulated and MYC is downregulated in neuroendocrine prostate cancer. A. Heatmap showing expression of the indicated genes in CRPC-adenocarcinoma (CRPC-Adeno) and neuroendocrine prostate cancer (CRPC-NE) patient samples from the Beltran cohort. B. Violin plots comparing expression of MYC family members (MYC, MYCL, MYCN) between CRPC-Adeno and CRPC-NE samples in the Beltran cohort. C. Heatmap of the indicated genes across patient groups from the ProAtlas dataset, including nonmalignant prostate tissue, adenocarcinoma with different Gleason score, and neuroendocrine prostate cancer. D. Box plots comparing expression of MYC family genes across nonmalignant samples, adenocarcinoma stratified by Gleason score, and NEPC in the ProAtlas dataset. E. Heatmap of selected lineage and MYC family genes across prostate cancer cell lines from the CTPC dataset. F. Box plots comparing MYC family gene expression across prostate cancer cell lines from the CTPC dataset. G. RT-qPCR validation of MYC family gene expression across representative prostate cancer cell lines. H. Western blot analysis of MYC family protein levels across prostate cancer cell lines.

Elevated MYCL expression coincided with increased expression of canonical neuroendocrine markers, including ASCL1, INSM1, and CHGA (Fig. 1A, C, E), whereas MYC expression was negatively correlated with neuroendocrine markers and positively associated with genes downregulated in NEPC, such as REST, YAP1, and KLK3. Notably, MYCL and neuroendocrine-associated markers were also expressed at relatively high levels in VCaP cells, an amphicrine PCa model, suggesting a potential role for MYCL in intermediate or transitional lineage states (Fig. 1E, F).

RT-qPCR and immunoblot analyses confirmed elevated MYCL mRNA and protein levels in NCI-H660 cells relative to adenocarcinoma models, while MYC mRNA was reduced and MYC protein remained undetectable (Fig. 1G, H). The apparent large fold-change in MYCN mRNA (Fig. 1G) primarily reflected its absence in other cell lines, as absolute MYCN transcript levels remained low (high Cq values), and MYCN protein was not detected in any model.

Together, these observations establish MYCL as a neuroendocrine-lineage-selective MYC family member in advanced PCa and reveal a reciprocal loss of MYC, consistent with a MYC family switch, distinguishing adenocarcinoma from NEPC during lineage transdifferentiation.

### MYCL impairs cell adhesion and induces cytoskeletal remodeling in prostate adenocarcinoma cells

MYCL overexpression in prostate adenocarcinoma cells did not markedly alter overall cell proliferation, as evidenced by comparable cell counts between control and MYCL-expressing C4-2B cells (Fig. 2A, B); similar results were observed in PC3 and TKO cell lines. In contrast, MYCL induction significantly impaired cell adhesion, with MYCL-overexpressing cells displaying reduced attachment and altered cellular morphology in adhesion assays (Fig. 2C). Consistent with these observations, live-cell imaging (sFig. 1B) revealed impaired cell spreading and sustained defects in attachment following seeding, characterized by rounded and poorly adherent MYCL-overexpressing cells.

**Fig. 2:**
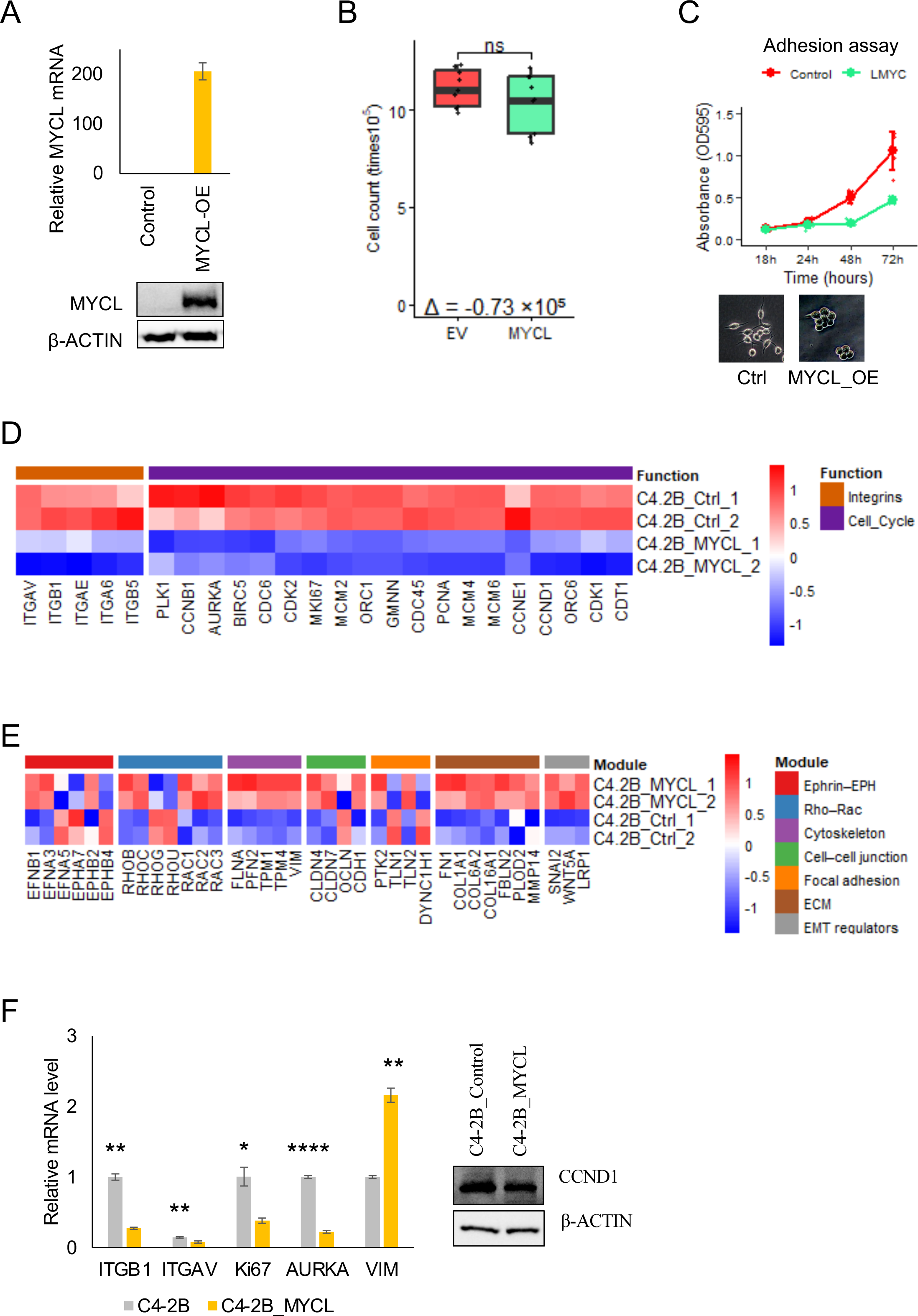
MYCL impairs cell adhesion and induces cytoskeletal remodeling in prostate adenocarcinoma cells. A. Confirmation of MYCL overexpression in C4-2B cells at both mRNA and protein levels. B. Cell proliferation assays showing comparable growth rates between control and MYCL-overexpressing prostate adenocarcinoma cells. C. Cell adhesion assays demonstrating reduced attachment of MYCL-overexpressing cells over time compared with controls. Representative phase-contrast images (bottom) illustrate altered cellular morphology and reduced adhesion in MYCL-overexpressing cells. D. Transcriptomic analysis revealing downregulation of integrin and cell-cycle associated genes. E. Heatmap of altered cytoskeletal and cell-cell adhesion genes, including Ephrin-EPH signaling, Rho-Rac GTPase signaling, focal adhesion, and extracellular matrix remodeling. F. RT-qPCR validation confirming reduced expression of integrins (ITGB1, ITGAV) and proliferation markers (KI67, AURKA), alongside increased expression of the mesenchymal marker VIM. Right: Western blot analysis showing reduced expression of CCND1 in MYCL-overexpressing cells. Statistical significance was determined using Student’s t-test; all comparisons shown were significant (P < 0.05).

Transcriptomic profiling further demonstrated that MYCL extensively altered the expression of adhesion- and cytoskeleton-associated gene networks. Integrin genes critical for extracellular matrix engagement-most notably ITGB1, a central mediator of integrin signaling were coordinately downregulated, together with reduced expression of multiple cell-cycle–associated transcripts, potentially reflecting impaired adhesion-dependent growth signaling (Fig. 2D). These transcriptional changes were accompanied by pronounced changes in the expression of cytoskeletal and adhesion-related genes, including Ephrin-EPH signaling, Rho-Rac GTPase pathways, focal adhesion, and extracellular matrix remodeling (Fig. 2E). Consistent with these findings, RT-qPCR validation confirmed reduced expression of integrins (ITGB1, ITGAV) and proliferation-associated markers (MKI67, AURKA), alongside increased expression of the mesenchymal marker VIM (Fig. 2F, left). Western blot analysis further demonstrated reduced CCND1 protein levels, consistent with attenuation of cell-cycle progression (Fig. 2F, right).

Together, these data indicate that MYCL drives cytoskeletal and cell-adhesion remodeling in prostate adenocarcinoma cells, with minimal effects on overall proliferative capacity.

### MYCL drives neuroendocrine-like transcriptional reprogramming and suppresses MYC signaling

Transcriptomic profiling revealed that MYCL overexpression induces broad transcriptional reprogramming. Gene set enrichment analysis of Hallmark pathways demonstrated enrichment of hypoxia, epithelial–mesenchymal transition, glycolysis, KRAS signaling, and apical surface pathways, accompanied by suppression of androgen responsive genes, MYC targets, and Notch signaling in MYCL-overexpressing cells (Fig. 3A). Gene Ontology analysis further revealed upregulation of multiple differentiation-associated biological processes, alongside coordinated downregulation of DNA replication, DNA repair, and chromosome organization pathways, indicating attenuation of proliferative and genome maintenance programs (Fig. 3B). Notably, neuronal and axon-related gene programs, including axon guidance, neuron projection development, regulation of neuronal differentiation, and nervous system processes were significantly enriched upon MYCL overexpression (Fig. 3C).

**Fig. 3:**
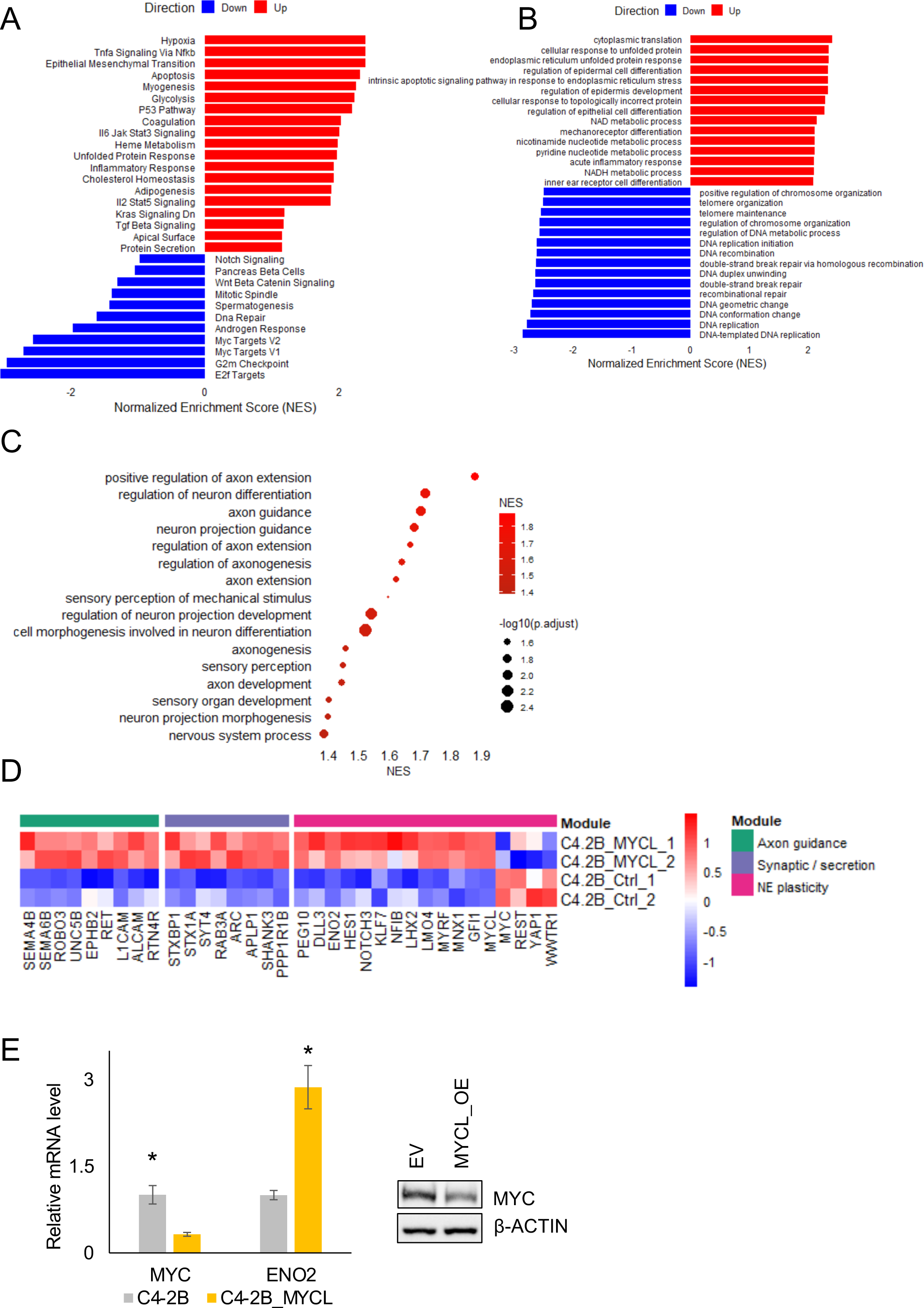
MYCL drives neuroendocrine-like transcriptional reprogramming and suppresses MYC signaling. A. Gene set enrichment analysis (GSEA) of Hallmark pathways showing enrichment of hypoxia, epithelial–mesenchymal transition, glycolysis, KRAS signaling, and apical surface pathways, with concurrent suppression of androgen response, MYC target genes, and Notch signaling in MYCL-overexpressing cells. B. GSEA of Gene Ontology biological processes revealing upregulation of differentiation-associated processes, alongside downregulation of DNA replication, DNA repair, and chromosome organization pathways upon MYCL overexpression. C. Dot plot highlighting enrichment of neuronal and axon-related gene programs, including axon guidance, neuron projection development, and nervous system processes, in MYCL-overexpressing cells. D. Heatmap of representative genes involved in axon guidance, synaptic/secretion pathways, and neuroendocrine (NE) plasticity, demonstrating coordinated upregulation of neuronal and NE-associated transcriptional modules in MYCL-overexpressing C4-2B cells compared with controls. E. RT-qPCR validation showing reduced MYC and increased ENO2 expression in MYCL-overexpressing cells (left), with corresponding Western blot analysis confirming suppression of MYC protein levels (right). Statistical significance was determined using Student’s t-test (P < 0.05).

Consistent with these findings, heatmap analysis demonstrated coordinated upregulation of genes associated with axon guidance, synaptic/secretion pathways, and neuroendocrine plasticity in MYCL-overexpressing C4-2B cells compared with controls (Fig. 3D). RT-qPCR validation confirmed reduced MYC expression together with increased expression of the neuroendocrine marker ENO2, and Western blot analysis corroborated suppression of MYC protein levels (Fig. 3E). Together, these findings support a MYC family switch in which MYCL expression coincides with repression of MYC signaling and activation of neuroendocrine-associated transcriptional programs. Collectively, these data indicate that MYCL drives transcriptional programs associated with neuronal differentiation and neuroendocrine plasticity while repressing androgen response and MYC-dependent signaling pathways in prostate adenocarcinoma cells.

### MYCL suppresses androgen receptor signaling and promotes resistance to AR pathway inhibition

MYCL overexpression was associated with marked suppression of androgen receptor (AR) signaling in prostate adenocarcinoma cells. Gene set enrichment analysis revealed significant downregulation of the Hallmark androgen response pathway in MYCL-overexpressing C4-2B cells compared with controls (Fig. 4A). Consistently, heatmap analysis of AR-regulated genes demonstrated coordinated repression of AR-induced genes alongside relative enrichment of AR-repressed genes that are typically induced following androgen deprivation (Fig. 4B). RT-qPCR analysis confirmed reduced expression of AR and canonical AR target genes, including KLK3, as well as luminal epithelial differentiation markers NKX3-1 and SPDEF, with corresponding reductions in AR and NKX3-1 protein levels validated by Western blotting (Fig. 4C).

**Fig. 4:**
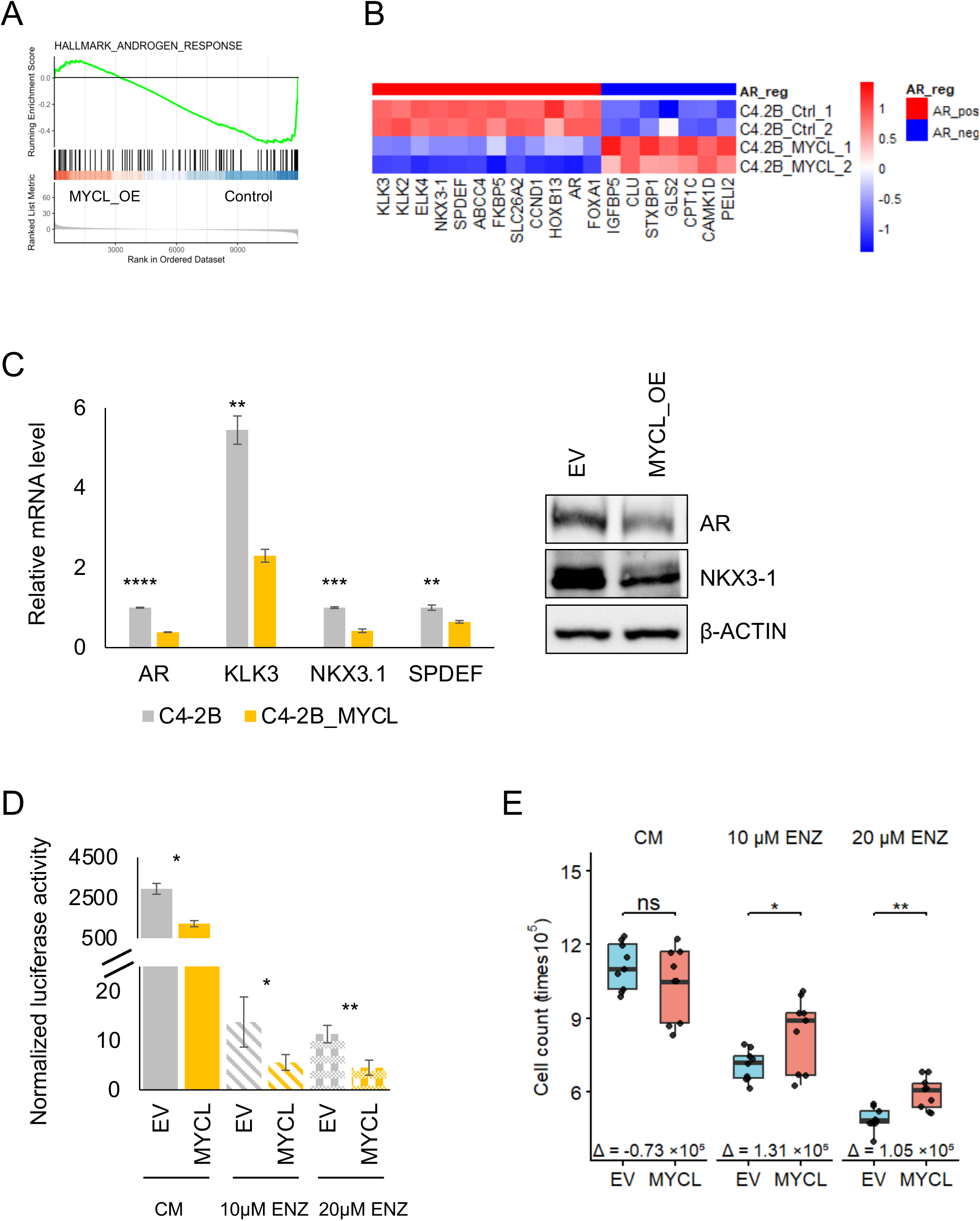
MYCL suppresses androgen receptor signaling and promotes resistance to AR pathway inhibition. **A.** Gene set enrichment analysis (GSEA) showing significant suppression of the Hallmark androgen response pathway in MYCL-overexpressing C4-2B cells compared with controls. **B.** Heatmap of AR-regulated genes in MYCL-overexpressing cells demonstrating coordinated downregulation of AR-positive targets and relative enrichment of AR-negative target genes that are typically induced following androgen deprivation. **C.** RT-qPCR analysis showing reduced expression of AR and canonical AR target genes, including KLK3 and luminal epithelial differentiation markers (NKX3-1 and SPDEF), in MYCL-overexpressing cells (left), with corresponding Western blot analysis confirming reduced AR and NKX3-1 protein levels (right). **D.** AR-responsive luciferase reporter assay demonstrating reduced AR transcriptional activity upon co-transfection with MYCL under control conditions and following enzalutamide (ENZ) treatment. **E.** Cell viability assays showing altered growth responses of MYCL-overexpressing cells compared with controls. Under complete media (CM), MYCL-overexpressing cells exhibit reduced proliferation, whereas under increasing concentrations of enzalutamide, MYCL-overexpressing cells display relatively improved proliferation compared with control cells. Statistical significance was determined using Student’s *t*-test (*P* < 0.05).

Functional assessment using an AR-responsive luciferase reporter assay, in which the reporter was co-transfected with MYCL, demonstrated significantly reduced AR transcriptional activity under basal conditions and following enzalutamide treatment (Fig. 4D). Consistent with attenuation of AR dependence, cell viability assays revealed reduced proliferation of MYCL-overexpressing cells in the absence of enzalutamide, whereas these cells exhibited relatively improved proliferation compared with control cells at increasing concentrations of enzalutamide (Fig. 4E). Together, these findings indicate that MYCL suppresses AR signaling and luminal differentiation programs, promoting a shift toward AR-independent growth states that may confer relative tolerance to AR pathway inhibition.

### MYCL is required to maintain neuroendocrine lineage identity in NEPC cells

To determine whether MYCL is required to maintain neuroendocrine (NE) lineage identity, MYCL was depleted in the NEPC cell line NCI-H660 using siRNA. Efficient knockdown of MYCL was confirmed at both the mRNA and protein levels (Fig. 5A). Global transcriptomic analysis revealed that loss of MYCL led to widespread reprogramming of gene expression rather than isolated pathway changes. Gene set enrichment analysis of Hallmark pathways demonstrated enrichment of MYC target gene programs and apical junction pathways following MYCL depletion, together with reduced enrichment of mTORC1 signaling, apical surface, glycolysis, and hypoxia pathways (Fig. 5B), suggesting a shift away from the metabolic and stress-adaptive programs characteristic of NEPC. Consistent with this, Gene Ontology analysis revealed marked downregulation of biological processes associated with neuronal fate commitment and neuroendocrine cell differentiation, alongside relative upregulation of DNA replication–related transcriptional programs (Fig. 5C).

**Fig. 5:**
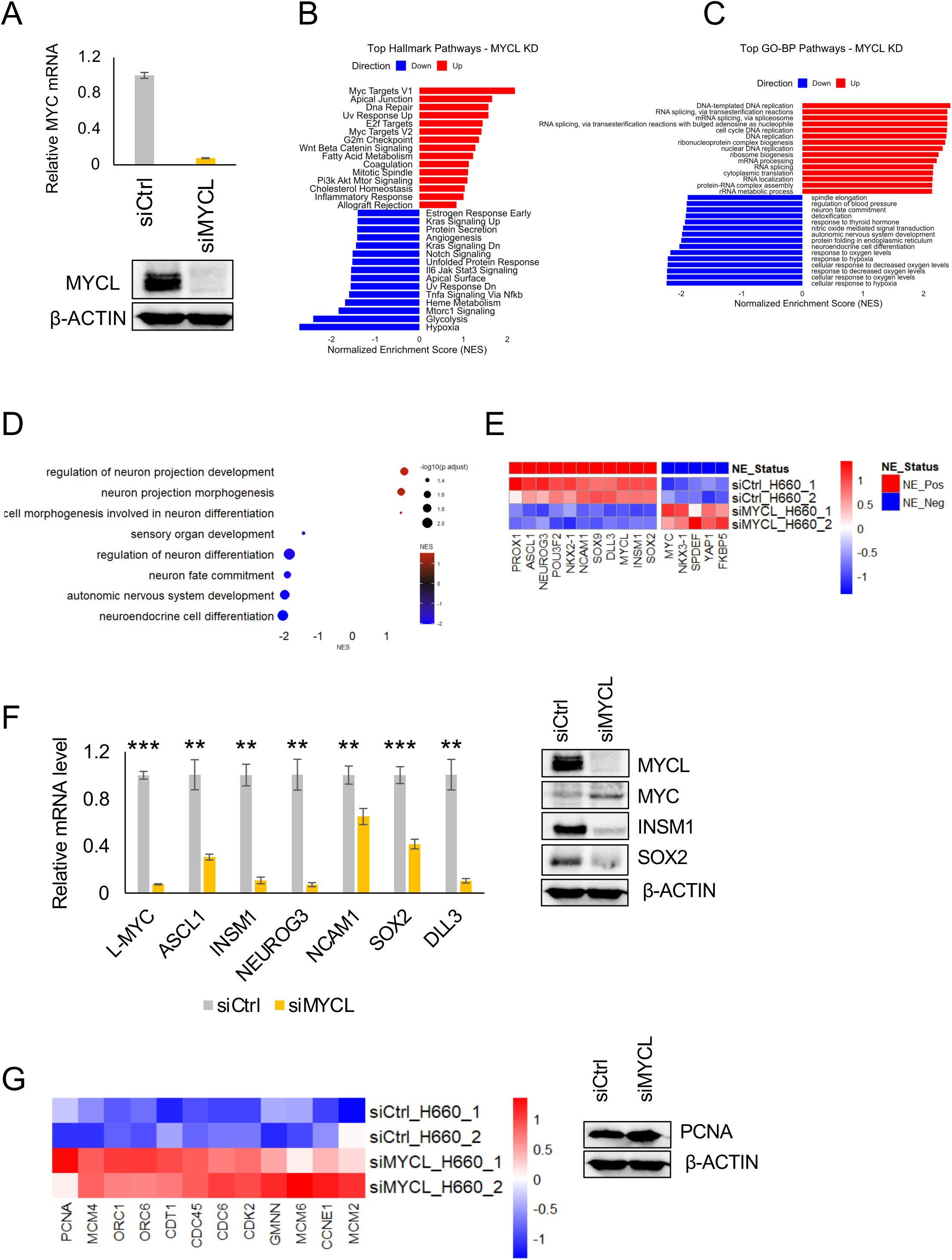
MYCL is required to maintain neuroendocrine lineage identity in NEPC cells. A. Efficient knockdown of MYCL in NCI-H660 cells confirmed by RT-qPCR (top) and Western blot analysis (bottom). B. Gene set enrichment analysis (GSEA) of Hallmark pathways following MYCL knockdown showing enrichment of MYC target gene programs and apical junction pathways, together with downregulation of mTORC1 signaling, apical surface, glycolysis, and hypoxia pathways. C. GSEA of Gene Ontology biological processes revealing downregulation of neuronal fate commitment and neuroendocrine cell differentiation pathways, alongside upregulation of DNA replication–related processes upon MYCL depletion. D. Dot plot highlighting negative enrichment of neuronal and neuroendocrine differentiation programs and positive enrichment of neuron morphogenesis–related pathways, including neuron projection development and neuron projection morphogenesis, following MYCL knockdown. E. Heatmap of lineage-associated markers demonstrating coordinated downregulation of neuroendocrine (NE) genes and concomitant upregulation of epithelial and adenocarcinoma-associated markers, including NKX3-1, SPDEF, FKBP5 (an AR target), and MYC, in MYCL-depleted H660 cells compared with controls. F. RT-qPCR (left) and Western blot (right) validation showing reduced expression of neuroendocrine markers, including ASCL1, INSM1, NEUROG3, NCAM1, SOX2, and DLL3, upon MYCL knockdown. G. Heatmap of cell-cycle–associated genes demonstrating reactivation of DNA replication and proliferation programs following MYCL depletion, with corresponding Western blot analysis confirming increased PCNA expression. Statistical significance was determined using Student’s t-test (P < 0.05).

Notably, dot plot analysis showed negative enrichment of neuronal and neuroendocrine differentiation pathways, while processes related to neuron morphogenesis and projection remodeling were relatively enriched (Fig. 5D), indicating disruption of mature NE lineage programs rather than global neuronal activation. This finding is consistent with the morphological alterations observed upon MYCL overexpression, suggesting that MYCL also contributes to regulation of cellular morphogenesis in NEPC. Heatmap analysis further demonstrated coordinated downregulation of canonical neuroendocrine lineage markers, accompanied by upregulation of epithelial and adenocarcinoma-associated genes, including NKX3-1, SPDEF, FKBP5, and MYC, which are typically suppressed in NEPC (Fig. 5E). These transcriptional changes were validated by RT-qPCR and Western blot analysis, confirming reduced expression of key neuroendocrine regulators ASCL1, INSM1, NEUROG3, NCAM1, SOX2, and DLL3 upon MYCL depletion (Fig. 5F). Collectively, these data indicate that MYCL is required to sustain neuroendocrine lineage identity in NEPC cells, and that loss of MYCL triggers coordinated transcriptional reprogramming away from a neuroendocrine state toward a more adenocarcinoma-like gene expression profile (Fig. 5G).

### Epigenetic and transcriptional mechanisms underlie MYCL upregulation in neuroendocrine prostate cancer

To determine whether elevated MYCL expression in NEPC is driven by genomic alterations, copy-number data from the Beltran cohort were analyzed. No MYCL amplification was detected in NEPC samples, indicating that MYCL overexpression is unlikely to be driven by gene dosage and is instead mediated by non-genetic mechanisms (Supplementary Fig. 2A).

To investigate epigenetic regulation, CCLE RRBS data were analyzed to assess CpG methylation at transcription start site (TSS) clusters across prostate cancer cell lines. The TSS regions of both MYCL and MYC were consistently hypomethylated across cell lines, suggesting a transcriptionally permissive epigenetic state (Fig. 6A). In contrast, the MYCN promoter exhibited overall hypermethylation, with the exception of 22Rv1 cells, consistent with epigenetic repression. These findings highlight differential methylation-based regulation among MYC family members and suggest that the MYCL locus is broadly accessible for transcription across PCa subtypes.

**Fig. 6:**
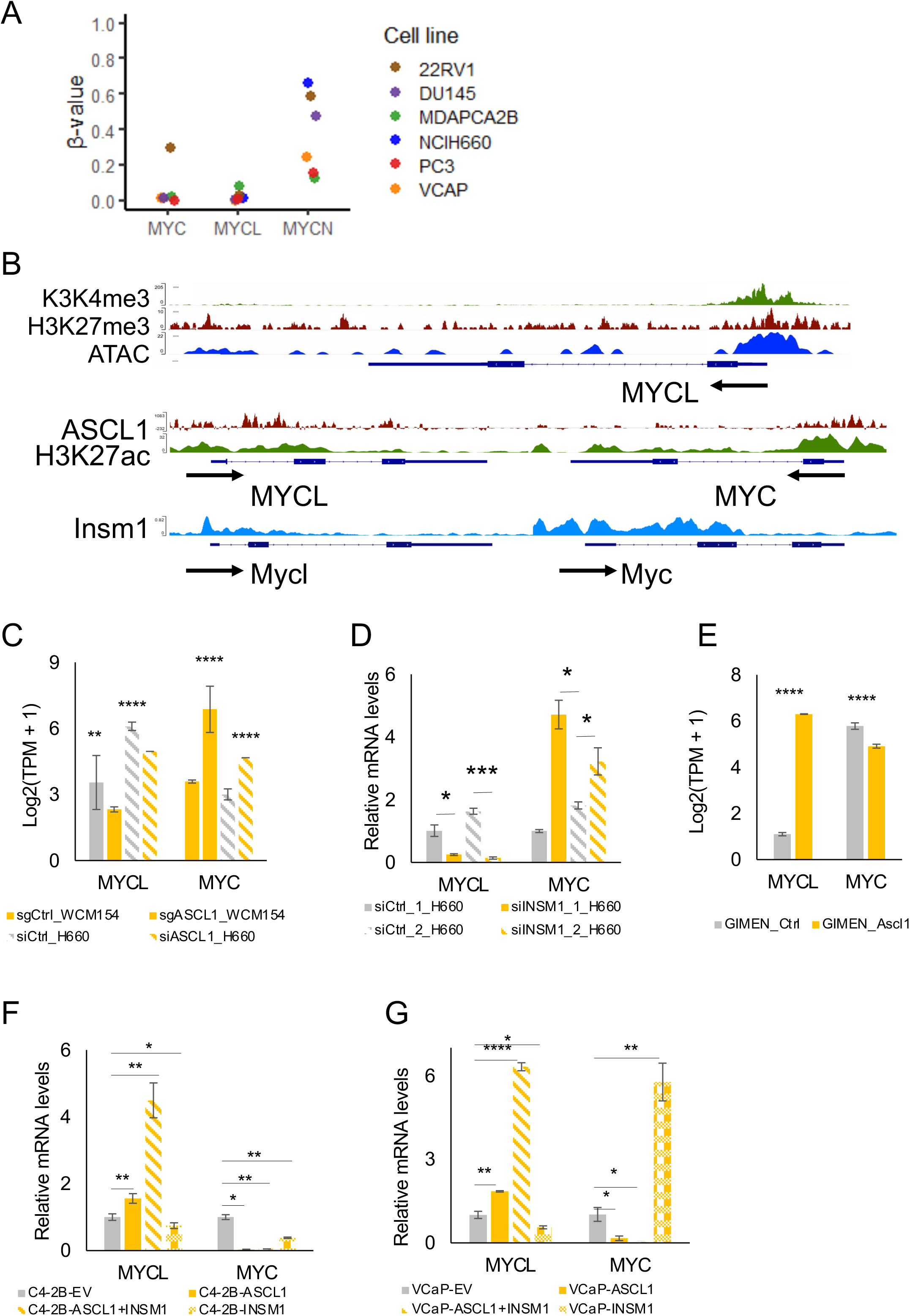
Conserved ASCL1-INSM1-MYCL regulatory axis in NEPC. A. CCLE RRBS analysis of CpG methylation at transcription start site (TSS) clusters across prostate cancer cell lines, showing relative hypomethylation of the MYCL and MYC promoters and hypermethylation of MYCN, indicating a permissive epigenetic state for MYCL and MYC transcription in these cells. B. ChIP-seq and ATAC-seq profiles showing chromatin accessibility and histone modifications (H3K4me3 and H3K27me3) at the MYCL and MYC loci in LNCaP cells, consistent with a bivalent, poised chromatin state. Middle panels show ASCL1 binding at the MYCL and MYC promoter regions together with H3K27ac enrichment in NCI-H660 cells. INSM1 binding at the MYCL and MYC loci is shown in mouse pancreatic β cells, where both MYCL and INSM1 are expressed, supporting conserved transcriptional regulation at these loci. C. Expression of MYCL and MYC following ASCL1 knockdown in NEPC patient-derived organoid (WCM154) and NCI-H660 cells, showing reduced MYCL expression with reciprocal changes in MYC. D. RT-qPCR analysis in NCI-H660 cells showing decreased MYCL and increased MYC expression following INSM1 knockdown, indicating coordinated regulation of MYCL within the ASCL1–INSM1 neuroendocrine transcriptional network. E. Expression of MYCL and MYC in neuroblastoma-derived GIMEN cells demonstrating induction of MYCL and suppression of MYC upon ASCL1 overexpression. F. RT-qPCR analysis in C4-2B cells showing induction of MYCL and repression of MYC following ASCL1 overexpression, with partial modulation by INSM1 co-expression. G. RT-qPCR analysis in VCaP cells showing robust induction of MYCL and suppression of MYC following ASCL1 overexpression, with modulation by INSM1. Statistical significance was determined using Student’s *t*-test (*P* < 0.05).

To further assess transcriptional accessibility, publicly available ATAC-seq and ChIP-seq datasets from LNCaP (adenocarcinoma) cells were examined. The MYCL promoter exhibited open chromatin together with a bivalent chromatin configuration, marked by co-occurrence of the activating H3K4me3 and repressive H3K27me3 histone modifications (Fig. 6B). Such bivalent states are characteristic of genes poised for activation during lineage transitions, suggesting that MYCL is epigenetically primed for transcriptional activation during neuroendocrine differentiation. ChIP-seq analysis further revealed ASCL1 binding at the MYCL promoter in NEPC cells (NCI-H660), as well as INSM1 binding at the MYCL locus in mouse pancreatic β cells-an endocrine cell type that co-expresses INSM1 and MYCL-supporting conserved transcriptional regulation. Notably, both ASCL1 and INSM1 also bound the MYC locus, suggesting coordinated and reciprocal regulation of MYCL and MYC by neuroendocrine lineage-defining transcription factors.

Functional validation in NEPC patient-derived organoids (WCM154) and NCI-H660 cells demonstrated that knockdown of ASCL1 or INSM1 resulted in reduced MYCL expression accompanied by increased MYC expression (Fig. 6C, D), indicating opposing regulation of these MYC family members within the neuroendocrine transcriptional network. Conversely, ASCL1 overexpression in neuroblastoma-derived GI-ME-N cells increased MYCL and decreased MYC expression (Fig. 6E). Similar induction of MYCL was observed following ASCL1 overexpression in C4-2B cells (Fig. 6F) and the amphicrine PCa cell line VCaP (Fig. 6G), indicating that this regulatory axis extends beyond strictly neuroendocrine contexts.

To examine whether this transcriptional regulation is conserved across additional neuroendocrine cancer models, RNA-seq data from lung cancer cell lines were analyzed. MYCL expression was significantly higher in small-cell lung cancer (SCLC) compared with non-small-cell lung cancer (NSCLC), consistent with its association with neuroendocrine identity (Supplementary Fig. 2B). MYCL expression strongly correlated with the neuroendocrine lineage transcription factors ASCL1 and INSM1 (Supplementary Fig. 2C). Consistently, RNA-seq analysis following ASCL1 knockdown in SCLC cell lines (NCI-H2107, NCI-H209, and DMS53) demonstrated significant downregulation of MYCL expression (Supplementary Fig. 2D).

Collectively, these findings demonstrate that MYCL overexpression in NEPC is not driven by genomic amplification but instead arises from a permissive epigenetic landscape coupled with transcriptional activation by a conserved ASCL1-INSM1 regulatory network. The reciprocal regulation of MYCL and MYC by neuroendocrine lineage transcription factors supports a dynamic MYC family regulatory switch that reinforces neuroendocrine identity across cancer models.

## DISCUSSION

Neuroendocrine prostate cancer (NEPC) represents a highly aggressive and treatment-resistant subtype of prostate cancer (PCa) that most commonly emerges through lineage plasticity from adenocarcinoma, particularly under the selective pressure imposed by androgen receptor pathway inhibitors (ARPIs). Unlike classical tumor progression driven by the acquisition of new oncogenic mutations, this transdifferentiation process is largely governed by epigenetic remodeling and reprogramming of transcriptional networks(Yamada & Beltran, 2021) (Yamada & Beltran, 2021)(Yamada & Beltran, 2021) (Yamada & Beltran, 2021). A defining hallmark of NEPC is the loss of AR signaling accompanied by the activation of neuroendocrine (NE) lineage genes, orchestrated by master regulators such as ASCL1 and INSM1. However, the downstream transcriptional effectors that translate these lineage cues into stable and self-reinforcing NE identity remain incompletely understood.

In this study, we identify MYCL as a previously unrecognized, lineage-specific member of the MYC family that is selectively upregulated in NEPC and is critical for both the establishment and maintenance of the neuroendocrine phenotype. In contrast to MYC, which is frequently overexpressed in AdPC and is closely linked to AR-dependent proliferative signaling, MYCL expression is selectively enriched in NEPC and shows strong concordance with canonical NE markers. This divergent expression pattern suggests a functional reprogramming of MYC family usage during lineage plasticity. Our findings support a model in which MYCL acts as a downstream effector of the NE lineage program, operating within an epigenetically permissive chromatin landscape and reinforced by NE master transcription factors. Rather than serving merely as a passive marker of neuroendocrine differentiation, MYCL likely functions as a lineage-stabilizing transcriptional amplifier that sustains NE gene expression programs while simultaneously suppressing AR-driven transcriptional states.

### A lineage-associated MYC family switch in prostate cancer progression

MYC family oncogenes have long been implicated in prostate cancer pathogenesis; however, their lineage-specific deployment during disease progression has remained poorly defined. MYC amplification and overexpression are hallmarks of adenocarcinoma prostate cancer (AdPC), where MYC drives proliferation, metabolic reprogramming, and androgen receptor (AR) signaling output. In contrast, MYCN amplification has been reported in a subset of neuroendocrine prostate cancer (NEPC) cases, where it cooperates with AKT signaling and epigenetic regulators such as EZH2 to suppress AR activity and promote neuroendocrine features. Our data uncover a distinct and complementary paradigm in which MYCL is selectively and robustly upregulated in NEPC, while MYC is concurrently suppressed and MYCN remains low at both transcript and protein levels in the majority of patient samples and experimental models examined.

This reciprocal MYCL-MYC expression pattern is consistently observed across independent patient cohorts, bulk and single-cell transcriptomic datasets, and prostate cancer cell line models. Notably, MYCL expression is detectable within AR-high adenocarcinoma subpopulations, suggesting that MYCL activation may precede or accompany early lineage transitions rather than representing a late consequence of fully established neuroendocrine differentiation. These findings support a model in which prostate cancer progression involves a dynamic reconfiguration of MYC family usage that closely tracks lineage identity and transcriptional state.

Functionally, this MYC family switch reflects divergent transcriptional programs associated with epithelial–luminal versus neuroendocrine lineages. MYC is highly expressed in AdPC and castration-resistant prostate cancer (CRPC) but is strongly repressed in NEPC. In contrast, MYCL is markedly upregulated in NEPC and correlates inversely with MYC expression and luminal epithelial genes such as REST and KLK3, while showing strong concordance with canonical neuroendocrine markers. Importantly, a similar MYCL-centered transcriptional axis has been described in small-cell lung cancer, where MYCL expression aligns with ASCL1-driven neuroendocrine subtypes, suggesting that MYCL-associated programs may represent a conserved regulatory mechanism across neuroendocrine malignancies.

### MYCL as a driver of cytoskeletal remodeling and neuroendocrine plasticity

Unlike MYC, whose oncogenic activity is tightly coupled to cell-cycle progression and proliferative output, MYCL overexpression in prostate adenocarcinoma cells exerted minimal effects on overall growth rates. Instead, MYCL profoundly remodeled cell adhesion, morphology, and cytoskeletal organization. Coordinated downregulation of integrins and focal adhesion components, together with altered expression of cell-cycle regulators, was accompanied by enrichment of Rho-Rac GTPase signaling, Ephrin–EPH pathways, and mesenchymal markers, indicating a primary role for MYCL in regulating cell–matrix interactions and cellular architecture. These phenotypic and transcriptional changes are particularly striking given the well-established link between loss of adhesion, cytoskeletal plasticity, and neuroendocrine differentiation across multiple cancer types. In neuroendocrine prostate cancer, cells characteristically adopt rounded morphologies and exhibit reduced substrate attachment. Our data suggest that MYCL directly contributes to these traits by reprogramming adhesion and cytoskeletal networks, thereby promoting lineage flexibility and facilitating adaptation to androgen receptor-depleted environments.

### MYCL suppresses AR signaling and MYC-dependent transcriptional programs

A defining feature of neuroendocrine prostate cancer (NEPC) is the loss of androgen receptor (AR) signaling and luminal epithelial identity. We demonstrate that MYCL overexpression is sufficient to suppress AR transcriptional output, downregulate canonical AR target genes, and attenuate AR-driven reporter activity. Consistent with this transcriptional rewiring, MYCL-expressing cells exhibit increased tolerance to AR pathway inhibition, supporting a functional shift toward AR-independent survival strategies.

Concomitant with AR suppression, MYCL induces broad repression of MYC-dependent transcriptional programs and reduces MYC expression at both the mRNA and protein levels. This finding reveals a functional antagonism between MYCL and MYC that extends beyond correlative lineage association. Rather than acting redundantly, MYCL appears to actively displace MYC-driven programs linked to proliferation, cell-cycle progression, and genome maintenance, while favoring differentiation-associated, neuronal, and stress-adaptive transcriptional states. This functional divergence provides a mechanistic basis for the MYC family switch observed during neuroendocrine transdifferentiation.

Notably, a similar division of labor among MYC family members has been described in small-cell lung cancer (SCLC), where MYCL promotes neuronal-like transcriptional programs but is insufficient on its own to fully establish ASCL1-positive neuroendocrine lineage identity. In that context, replacement of MYC with MYCL shifts the transcriptional landscape toward neuroendocrine gene expression, particularly when MYC activity is diminished, and is accompanied by reconfiguration of chromatin accessibility at neuronal regulatory elements (Patel et al., 2021)(Patel et al., 2021)(Patel et al., 2021)(Patel et al., 2021). Together, these observations highlight a context-dependent role for MYCL-distinct from canonical MYC-in suppressing proliferative, lineage-incompatible programs while reinforcing neuroendocrine identity.

### MYCL is required to maintain neuroendocrine lineage identity

Loss-of-function studies in the NEPC cell line NCI-H660 demonstrate that MYCL is not only sufficient to induce neuroendocrine-like programs but is also required to maintain them. MYCL depletion resulted in coordinated downregulation of canonical neuroendocrine markers, including ASCL1, INSM1, SOX2, and DLL3, alongside reactivation of epithelial and adenocarcinoma-associated genes such as NKX3-1, SPDEF, FKBP5, and MYC. This transcriptional reversion was accompanied by reactivation of DNA replication and proliferation-associated programs, suggesting that MYCL actively enforces a differentiated, AR-independent neuroendocrine state.

Notably, MYCL knockdown did not simply abolish neuronal gene expression but instead disrupted mature neuroendocrine lineage programs while disturbing morphogenesis-related pathways. This distinction highlights MYCL’s role in stabilizing lineage identity, consistent with its function as a lineage maintenance factor.

### Epigenetic priming and conserved ASCL1–INSM1 regulation of MYCL

A key insight from this study is that MYCL activation in neuroendocrine prostate cancer (NEPC) is not driven by genomic amplification. Unlike MYC, which is frequently upregulated through copy number gain in adenocarcinoma prostate cancer (AdPC) (Qiu et al., 2022b)(Qiu et al., 2022b)(Qiu et al., 2022b)(Qiu et al., 2022b), the MYCL locus resides within an epigenetically permissive chromatin landscape characterized by promoter hypomethylation and bivalent chromatin marks. Such configurations are characteristic of lineage-specifying genes in developmental systems, enabling rapid transcriptional activation during differentiation and lineage reprogramming (Jeon & Tucker-Kellogg, 2020)(Jeon & Tucker-Kellogg, 2020)(Jeon & Tucker-Kellogg, 2020)(Jeon & Tucker-Kellogg, 2020). This epigenetic priming likely facilitates MYCL induction during the transition to NEPC.

We further identify ASCL1 and INSM1-core neuroendocrine lineage transcription factors-as direct upstream regulators of MYCL. ChIP-seq evidence demonstrates ASCL1 and INSM1 binding at the MYCL locus, and functional perturbation experiments across prostate and lung neuroendocrine models reveal that knockdown of either factor reduces MYCL expression while reciprocally increasing MYC levels. This coordinated regulation establishes a conserved ASCL1-INSM1-MYCL axis that provides a mechanistic framework for the MYC family switch observed during neuroendocrine transdifferentiation. In this model, MYCL functions downstream of canonical neuroendocrine transcriptional hierarchies, reinforcing lineage-specific gene expression while suppressing MYC-dependent programs.

Importantly, this regulatory architecture is conserved across neuroendocrine malignancies. In small-cell lung cancer (SCLC), MYCL is selectively expressed relative to non-small-cell lung cancer and is strongly associated with ASCL1- and INSM1-positive neuroendocrine subtypes. Prior studies have demonstrated mutually exclusive MYC family amplifications in SCLC and shown that MYC, MYCN, and MYCL drive distinct, lineage-specific transcriptional programs, with additional work revealing that these MYC family members operate through differential enhancer usage (Patel et al., 2021; Plotnik et al., 2024). In this context, MYCL is linked to neuronal and neuroendocrine gene expression, whereas MYC is associated with metabolic and non-neuroendocrine programs. Together, these findings support MYC-to-MYCL switching as a conserved feature of neuroendocrine reprogramming that stabilizes lineage commitment across tumor types.

### Biological and therapeutic implications

Although MYCL amplification is not a prominent feature in the Beltran NEPCa cohort, prior studies have reported focal MYCL amplification in clinically localized prostate tumors (Boutros et al., 2015). These events were more frequent than MYC amplification and mutually exclusive with other MYC family amplifications. Importantly, MYCL expression remained low in these tumors, suggesting epigenetic silencing in AdPC that may be unlocked during lineage reprogramming.

Interestingly, MYCL has also been shown to promote iPSC reprogramming more efficiently than MYC and MYCN, without inducing tumor formation (Nakagawa et al., 2010)(Nakagawa et al., 2010)(Nakagawa et al., 2010)(Nakagawa et al., 2010). This separation of reprogramming capacity from oncogenic transformation supports the idea that MYCL functions as a lineage-specifying factor rather than a classical oncogene. In NEPC, its upregulation in NEPC may reflect its association with transcriptional rather than proliferative signatures.

Given the challenges of directly targeting MYC proteins, upstream regulators of MYCL, such as ASCL1 and INSM1, or epigenetic mechanisms enabling its expression may provide novel therapeutic opportunities. In SCLC models, MYCL silencing reduced tumor development (Kim et al., 2016)(Kim et al., 2016)(Kim et al., 2016)(Kim et al., 2016), supporting its functional relevance and therapeutic potential in NE malignancies.

## CONCLUSIONS

This study identifies MYCL as a lineage-specific transcriptional factor selectively upregulated in NEPC. Unlike MYC, which is enriched in adenocarcinoma and linked to proliferation, MYCL is epigenetically upregulated and reinforces the NE phenotype through a conserved transcriptional program driven by transcriptional program by master lineage factors ASCL1 and INSM1. Our findings reveal a conserved MYC family switch that underpins NE transdifferentiation and establish MYCL as a key effector of lineage identity. These insights open new avenues for understanding and targeting lineage plasticity in NEPC and related malignancies.

## MATERIALS AND METHODS

### Bioinformatic analysis

To investigate the expression of MYC family genes and NE markers across different prostate cancer states, RNA-seq and copy number variation data from patients in the Beltran and SU2C cohorts were obtained from cBioPortal (Cerami et al., 2012; Gao et al., 2013)(Cerami et al., 2012; Gao et al., 2013)(Cerami et al., 2012; Gao et al., 2013)(Cerami et al., 2012; Gao et al., 2013). Expression and methylation data for prostate and expression data for lung cancer cell lines were obtained from the Cancer Cell Line Encyclopedia (CCLE) (Ghandi et al., 2019)(Ghandi et al., 2019)(Ghandi et al., 2019)(Ghandi et al., 2019) and included for comparative analysis. Normalized gene expression values were log₂-transformed prior to heatmap visualization using the R pheatmap package. Boxplots illustrating MYC family gene expression across pathological subtypes were generated using the ggplot2 package, and statistical differences between groups were assessed using the Wilcoxon rank-sum test.

Spearman correlation between MYC family genes and neuroendocrine (NE) lineage markers was performed for the Beltran and SU2C prostate cancer cohorts and prostate cancer cell lines using the Hmisc package in R. Correlation matrices were visualized as clustered heatmaps using pheatmap. For lung cancer cell lines, Spearman correlation was performed between MYCL expression and epigenetic regulators in MYCL-amplification-free lines from the CCLE dataset. MYCL-associated genes identified in lung cancer were then analyzed in the Beltran NEPC dataset to assess the preservation of MYCL-linked transcriptional networks. Dot heatmaps were generated using log₂-transformed CCLE RNA-seq data to visualize MYC family gene expression across prostate cancer cell lines, with dot size and color representing expression magnitude. For the Beltran cohort, log₂-transformed MYCL copy number and expression values were visualized as scatter plots with statistical annotations using the ggplot2 and ggpubr packages. Methylation β-values for MYC family genes in PCa cell lines were visualized using the pheatmap package, with a color gradient indicating a continuum from low (hypomethylated) to high (hypermethylated) states.

### Chip-Seq analysis

Epigenomic datasets were retrieved from GEO, including H3K27me3 and H3K4me3 ChIP-seq profiles in LNCaP cells from GSE148935 (Li et al., 2022)(Li et al., 2022)(Li et al., 2022)(Li et al., 2022) and ATAC-seq data in LNCaP cells from GSE139099 (Grubert et al., 2020)(Grubert et al., 2020)(Grubert et al., 2020)(Grubert et al., 2020), INSM1 ChIP-seq data in pancreatic β-cells were obtained from GSE54046 (Jia et al., 2015)(Jia et al., 2015)(Jia et al., 2015)(Jia et al., 2015) and ASCL1 ChIP-seq data in H660 cells were retrieved from GSE183198 (Nouruzi et al., 2022)(Nouruzi et al., 2022)(Nouruzi et al., 2022)(Nouruzi et al., 2022). H3K27ac ChIP-seq data in H660 cells were downloaded from GSE224421 (Tabrizian et al., 2023)(Tabrizian et al., 2023)(Tabrizian et al., 2023)(Tabrizian et al., 2023). All downloaded BigWig files were visualized using the Integrative Genomics Viewer (IGV) (Robinson et al., 2011)(Robinson et al., 2011)(Robinson et al., 2011)(Robinson et al., 2011). Except for INSM1 (mm10) and H660 ASCL1 and H3K27ac datasets (hg38), all data were aligned to the hg19 genome build.

### RNA-seq analysis of ASCL1 and MYC family perturbation models

RNA-seq datasets for ASCL1 knockdown in H660, H2107, H209 and DMS53 cells were obtained from the GEO datasets under accessions GSE183199 (Nouruzi et al., 2022)(Nouruzi et al., 2022)(Nouruzi et al., 2022)(Nouruzi et al., 2022), GSE151002 (Pozo et al., 2021)(Pozo et al., 2021)(Pozo et al., 2021)(Pozo et al., 2021), GSE129340 (Hokari et al., 2020)(Hokari et al., 2020)(Hokari et al., 2020)(Hokari et al., 2020) and, GSE179071 (Costanzo et al., 2022)(Costanzo et al., 2022)(Costanzo et al., 2022)(Costanzo et al., 2022) respectively. ASCL1 overexpression in GI-ME-N neuroblastoma cells was analyzed using dataset GSE214796 (Wang et al., 2023)(Wang et al., 2023)(Wang et al., 2023)(Wang et al., 2023).

Raw RNA-seq count data were processed in R. Gene lengths were retrieved from Ensembl using the biomaRt package, and transcripts per million (TPM) values were calculated by normalizing read counts to gene length and sequencing depth. TPM was computed by first calculating reads per kilobase (RPK), followed by scaling RPK values so that the sum per sample equals one million.

Differential expression analysis was performed using the DESeq2 package in R. Raw count data were used to construct a DESeqDataSet object, and genes with low expression (row sums ≤ 1) were filtered out. The DESeq2 pipeline was applied to estimate log₂ fold changes and adjusted p-values using the Benjamini–Hochberg correction, comparing experimental and control conditions.

A volcano plot was generated using ggplot2 and ggrepel to visualize differentially expressed genes. Genes were categorized as upregulated, downregulated, or not significant based on an adjusted p-value threshold of 0.05 and the direction of the log₂ fold change. MYC and MYCL genes were labeled on the plot if they were identified as differentially expressed.

Expression levels from biological replicates in each experimental condition were log₂-transformed as log₂(TPM + 1). For each gene and condition, mean expression and standard deviation were calculated. Bar plots displaying mean expression with error bars representing ± standard deviation were generated using ggplot2, with individual replicate data points overlaid as jittered dots. Log₂ fold changes and adjusted p-values from the differential expression analysis were annotated above the bars to indicate statistical significance.

### Cell culture

Prostate cancer cells were cultured in RPMI-1640 medium supplemented with 10% fetal bovine serum (FBS) and 1% penicillin-streptomycin at 37 °C in a humidified incubator with 5% CO₂. Cells were passaged upon reaching ∼80% confluency using 0.25% trypsin-EDTA and reseeded at 50% confluency for further experiments. HEK293A cells were cultured in Dulbecco’s Modified Eagle Medium (DMEM) supplemented with 10% FBS and 1% penicillin-streptomycin under the same incubation conditions.

### RT-qPCR and Western blot analysis

Total RNA was isolated from cells using the Zymo RNA isolation kit, following the manufacturer’s instructions. cDNA was synthesized using ABScript III Reverse Transcriptase (Abclonal). Quantitative PCR (qPCR) was performed using SYBR Green and gene-specific primers, following the two-step qPCR protocol provided by Abclonal. Ct values were analyzed using the ΔΔCt method to determine relative gene expression levels. Statistical analysis of RT-qPCR results was performed using an unpaired two-tailed Student’s t-test on ΔΔCt values, comparing control and experimental groups. A p-value < 0.05 was considered statistically significant. For western blot analysis cells were lysed in Laemmli SDS sample buffer and subjected to SDS-PAGE, followed by Western blotting. Primary antibodies used were β-actin (sc-47778, 1:500 dilution; Santa Cruz Biotechnology, Dallas, TX) and MYCL (#76266, 1:500 dilution; Cell Signaling Technology). An HRP-conjugated secondary antibody was used for detection. Protein bands were visualized using Thermo Scientific Pierce™ ECL Western Blotting Substrate and imaged with the ChemiDoc™ Touch Imaging System (Bio-Rad).

### Declaration of generative AI and AI-assisted technologies in the writing process

During the preparation of this work the authors used ChatGPT to improve readability. After using this tool, the authors reviewed and edited the content as needed and take full responsibility for the content of the publication.

## Supporting information

sFig 1 and sFig 2

## ACKNOWLEDGEMENTS

This research was supported by NIH R01 CA226285 and LSU Health Shreveport FWCC and Office of Research funding to XY.

