## Supplementary material for "A MYC Family Switch: L-MYC Drives and Maintains Neuroendocrine Lineage Programs in Prostate Cancer": sFig 1 and sFig 2

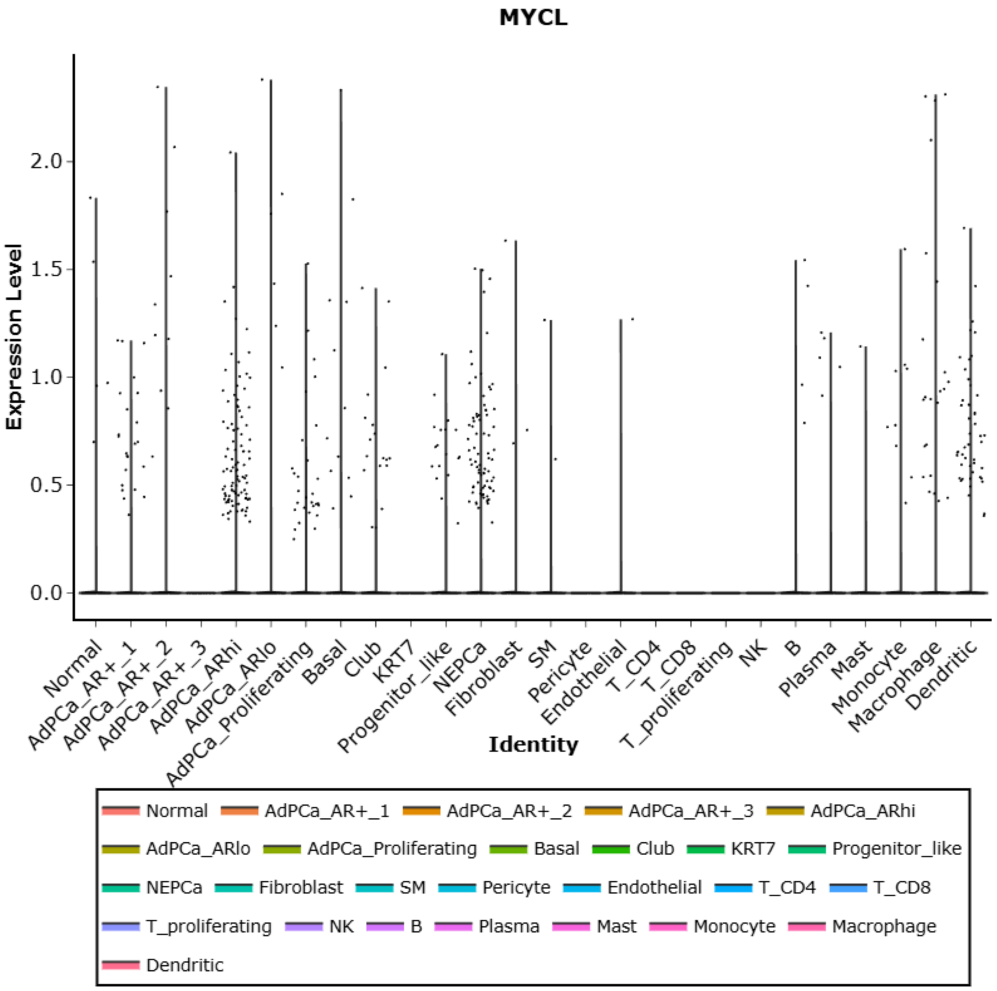


sFig. 1 Single-cell RNA-sequencing analysis showing MYCL expression across prostate epithelial and stromal cell populations, with elevated expression in neuroendocrine prostate cancer (NEPC) cells and AdPC_ARhi populations compared with other adenocarcinoma and non-epithelial cell types.

A

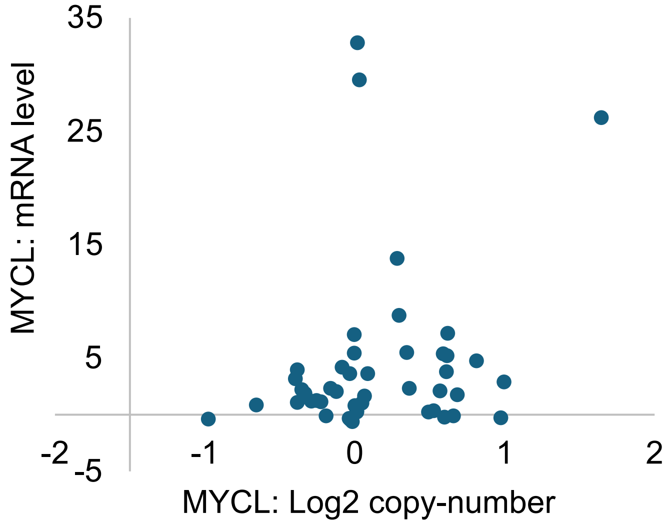


sFig. 2A Scatter plot showing the relationship between MYCL copy number (log2) and MYC mRNA expression across Beltran tumor samples, illustrating the absence of a strong positive correlation between MYCL genomic gain and MYC transcript levels.

B

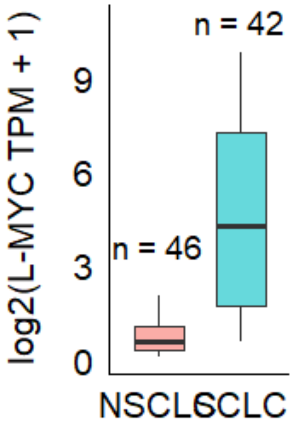


sFig. 2BBox plot comparing L-MYC expression [log2(TPM + 1)] between non-small cell lung cancer (NSCLC; *n* = 46) and small cell lung cancer (SCLC; *n* = 42) samples, demonstrating significantly higher L-MYC expression in SCLC.

C

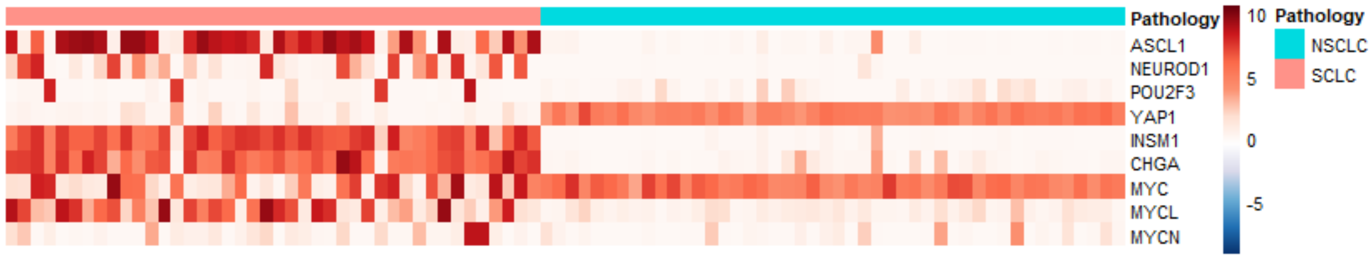


sFig. 2C Heatmap of lineage markers and MYC family gene expression across NSCLC and SCLC samples, showing enrichment of neuroendocrine markers (ASCL1, NEUROD1, INSM1, CHGA) together with MYCL in SCLC, and relative enrichment of MYC and YAP1 in NSCLC.

D

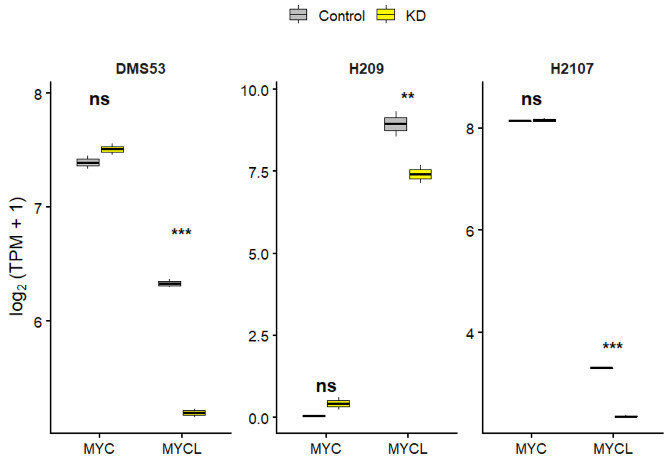


sFig. 2D Box plots showing expression of MYC and MYCL [log2(TPM + 1)] following ASCL1 knockdown (KD) compared with control in SCLC cell lines DMS53, H209, and H2107, indicating that ASCL1 regulates MYCL expression across models. Statistical significance was determined using DESeq2-adjusted P-values (padj; ns, not significant; padj < 0.05).
